# Enzymatic degradation of biofilm by metalloprotease from *Microbacterium* sp. SKS10

**DOI:** 10.1101/655092

**Authors:** Sandeep Kaur Saggu, Gopaljee Jha, Prakash Chandra Mishra

## Abstract

Enzymes have replaced or decreased usage of toxic chemicals for industrial and medical applications leading towards sustainable chemistry. In this study, we report purification and characterization of a biofilm degrading protease secreted by *Microbacterium* sp. SKS10. The protease was identified as a metalloprotease, Peptidase M16 using mass spectrometry. It showed optimum activity at 60 °C, pH 12 and retained its activity in the presence of various salts and organic solvents. The enzyme was able to degrade biofilms efficiently at enzyme concentration lower than other known enzymes such as papain, trypsin and □-amylase. The presence of this protease increased the accessibility of antibiotics inside the biofilm, and was found to be non-cytotoxic towards human epidermoid carcinoma cells (A431) at the effective concentration for biofilm degradation. Thus, this protease may serve as an effective tool for management of biofilms.

## 1 Introduction

Biofilms are a city of microbes (Watnick and Kolter, 2000) that adhere to each other forming complex tertiary structures that stick to a surface (Lopez et al., 2010). The microbial cells are embedded into a self-produced extracellular matrix comprising of polymeric substances such as polysaccharides, proteins, lipids and DNA (Jamal et al., 2015). The cells in biofilms have the capability to mutate and exchange genetic information helping them to survive in severe growth conditions (Singh et al., 2017). They are provided with various benefits such as physical defense from the host immune system and antimicrobials, sorption and storage of nutrients, tolerance to desiccation, and synchronization of virulence factor expression through quorum sensing. Oxygen deficiency and low metabolic activity make bacteria more tolerant to antibiotics (Singh et al., 2017). Rapid changes in pH among layers in biofilms may lead to the aggregation of organic acids and deactivation of the penetrating antimicrobial agents. *Staphylococcus aureus* (*S. aureus*) is one of the leading species in biofilm-associated infections which represent 80% of nosocomial infections. The main component of the *S. aureus* biofilm matrix is polysaccharide intercellular adhesin composed of β-1,6-linkedN-acetylglucosamine polymer and many proteins such as protein A, fibrinogen-binding proteins, clumping factor B etc. (Lister and Horswill, 2014).

Various chemicals such as phenols, alcohols, heavy metals and surfactants have been historically used for removal of biofilms (Araujo et al., 2011). Triclosan is one of the bisphenol compounds that have been used to degrade biofilms but this chemical possesses several health hazards such as disruption of mitochondrial function and calcium signaling, and alteration of homeostasis and immunological parameters (Weatherly and Gosse, 2017). Heavy metals such as mercury, silver, copper, zinc and nickel as well as chlorinated compounds have been used for biofilm dispersal but these chemicals are no longer in use due to their serious health hazards (Teitzel and Parsek, 2003). Due to various limitations of chemicals used for biofilm removal, there is a need to replace chemicals with enzymes.

The enzymes such as glycosidases, proteases and DNases have the potential to degrade the extracellular matrix (Algburi et al., 2017) resulting in release of planktonic cells and its constituents which are more easily accessible to antimicrobials (Fleming and Rumbaugh, 2017). As proteins are one of the major constituents of biofilms, the proteases are considered to be the most potential enzymes for biofilm removal (Lister and Horswill, 2014). Proteases such as aureolysin, proteinase K, spl protease, staphopain A and B produced by staphylococcal strains help in degradation of biofilms (Fleming and Rumbaugh, 2017). LapG secreted by *Pseudomonas putida* triggers the dispersal of biofilms through alteration of exopolysaccharide-binding protein, LapA (Gjermansen et al., 2010).

The present study deals with the purification and characterization of a protease from *Microbacterium* sp. SKS10 that has the potential for biofilm removal. We also report the isolation and characterization of this strain using morphological, biochemical and molecular methods. This study explores the potential of the protease in degrading biofilms and determines its toxicity on cultured human cells.

## 2 Materials and methods

### 2.1 Isolation and screening of microorganism

The bacterial strain SKS10 was isolated from soil of the Indian west coast region i.e. Mumbai, Maharashtra, (19.076 N, 72.877 E) and screened for its ability to hydrolyze casein protein present in skim milk agar plates (pH 10) [*peptone* (0.1%), *sodium chloride* (0.5%), *agar* (2%) and *skim milk* (10%)] incubated at 37 °C using serial dilution method for 48 hours. The colonies of bacteria forming a clear zone on these plates were isolated. This pure culture was maintained by subculturing on nutrient agar plates at 37 °C and stored as glycerol stocks at −80 °C.

### 2.2 Identification of strain SKS10

*Microbacterium sp.* SKS10 was characterized by morphological, biochemical and molecular methods. The morphological characteristics of a bacterial colony were observed using a stereomicroscope (Magnus, India). Cell morphology was studied by gram staining method using a brightfield microscope at 40x magnification (Nikon H600L, Japan) and scanning electron microscopy at a magnification of 10,000x (Zeiss Evo LS10, Germany).

The biochemical characteristics were evaluated by carbohydrate utilization tests using HiCarbo kit (HiMedia, India). The capability of strain for secretion of amylase, lipase, lysine decarboxylase, indole and acetylmethylcarbinol was determined by using respective media (Sivanandhini et al., 2015, Pammi et al., 2015) as per the manufacturer’s instructions.

A set of universal bacterial primers for 16S rRNA gene i.e. 8F (5’-AGAGTTTGATCCTGGCTCAG-3’) and 1492R (5’-GGTTACCTTGTTACGACTT-3’) was used to amplify 16S ribosomal DNA gene of the bacterial isolate by polymerase chain reaction (PCR) for molecular characterization. The gene sequence (sequenced by First Base Laboratories Sdn Bhd, Malaysia) was submitted in GenBank with accession number KY230497. Basic local alignment search tool (BLAST) was used to search GenBank (Johnson et al., 2008), ribosomal database project-II (RDPII) (Maidak et al., 2000) and EzTaxon (Chun et al., 2007) using 16S rRNA gene sequence. The 16S rRNA sequence of the isolated strain showed 100% identity with *Microbacterium paraoxydans*. Mega 7 software was used to build a phylogenetic tree with bootstrap value of 500 replicates by neighbor-joining method using maximum composite likelihood (Kumar et al., 2016). For this analysis, homologous sequences of 16S rRNA gene of *Microbacterium paraoxydans* were retrieved from RDPII database where *Actinocatenispora thailandica* was selected as an out group.

### 2.3 Protease assay and protein estimation

Protease activity was measured by modified Lowry method (Boominadhan et al., 2009; Lowry et al., 1951) whereas protein concentration was estimated by Bradford method (Bradford 1976). Reaction mixture containing 1% casein (Sisco Research Laboratories Pvt. Ltd., India, www.srlchem.com) as a substrate (400 μl) in 0.1M potassium chloride–sodium hydroxide (pH 12) and purified enzyme (200 μl) was incubated for half an hour at 37 °C. The reaction was terminated by addition of 600 μl of 10% trichloroacetic acid (Sisco Research Laboratories Pvt. Ltd., India, www.srlchem.com) followed by incubation for 10 minutes at room temperature. The reaction mixture was centrifuged at 10,000 g. Supernatant (200 μl) was added to 0.4M sodium carbonate (800 μl) and 1N Folin phenol Ciocalteau reagent (50 µl), incubated for half an hour at room temperature, and the absorbance measured at 660 nm. One unit enzyme activity was defined as the amount of enzyme that is needed to release 1 µg tyrosine per ml per minute under standard assay conditions.

### 2.4 Protease production and purification

*Microbacterium* sp. SKS10 was grown in 50 ml of nutrient broth in an Erlenmeyer flask (250 ml). The protease production was done using 5% inoculum of seed culture in Erlenmeyer flask (1L) having 2% casein in nutrient broth (pH 8) at 30 °C with agitation of 150 revolutions per minute for 48 hours.

Protein was precipitated from supernatant obtained after centrifugation following 70% ammonium sulphate fractionation (Sisco Research Laboratories Pvt. Ltd., India, www.srlchem.com). The precipitated protein was recovered after centrifugation at 10,000 g for 15 minutes and dissolved in minimal volume of buffer (pH 8) containing 50 mM Tris and 300 mM sodium chloride. This precipitated partially purified enzyme was further purified by gel permeation chromatography on a Superdex 75 10/300 G column (30 cm*10 mm) at a flow rate of 0.5 ml/min using GE AKTA Prime. The purity of elutes was checked by sodium dodecyl sulphate-polyacrylamide gel electrophoresis (SDS-PAGE) using silver staining method (Sambrook et al., 2001). 1 mg/ml chymotrypsin (25 kDa) was run as a standard on the same gel permeation chromatography column. The purified fractions obtained after gel permeation chromatography were pooled, stored at −20 °C and used for protease characterization. Zymogram analysis was used to check protease activity (Section 2.5). Specific activity was calculated as enzyme activity per mg protein at each step of purification.

### 2.5 Zymogram analysis

The activity of proteases in gel was detected using casein zymography (Bester et al., 2010). After electrophoresis, SDS-PAGE containing 0.1% casein was soaked in 2.5% (v/v) Triton X-100 at room temperature for 1 hour followed by overnight incubation in 50 mM Tris buffer (pH 8). This gel was stained with Coomassie Brilliant Blue-R250. The appearance of a zone of clearance was considered as the existence of protease activity. SDS-PAGE as well as zymography was performed under reducing and non-reducing conditions to investigate the role of disulphide bonds in protease activity.

### 2.6 Protease characterization

To study thermostability of the enzyme, the enzyme was preheated for 30 minutes at different temperatures (20 °C-80 °C) followed by protease activity under standard assay conditions. The time period up to which the protease was thermostable was determined by incubating the protease at 60 °C for different time intervals from 30-180 minutes (30, 60, 90, 120, 150 and 180 minutes) before measuring protease activity. The effect of pH on enzyme activity was determined by varying the pH of the assay mixture using various buffer systems at 0.1 M concentration under standard assay conditions. Various buffers used were: potassium phosphate buffer (pH 6, 7 and 8), carbonate–bicarbonate buffer (pH 9 and 10), sodium bicarbonate–sodium hydroxide buffer (pH 11) and potassium chloride–sodium hydroxide (pH 12). The optimum temperature for protease activity was evaluated by incubating the reaction mixture at optimum pH (pH 12) for 30 minutes at different temperatures (20 °C-80 °C).

The effect of metal salts on the enzyme activity was investigated by incubating the reaction mixture with chloride salts of various metal ions (Na^+^, Li^+^, Mg^2+^, K^+^, Ba^2+^, Ca^2+^, Mn^2+^, Cd^2+^, Zn^2+^, Fe^3+^, and Co^2+^) at 10 mM concentration for 30 minutes. An equal volume of water was added instead of metal salts in the control reaction. To examine the effect of various solvents on protease activity, the reaction mixture was incubated with various solvents (ethyl acetate, formamide, carbon tetrachloride, methanol, ethylene glycol, dodecane, hexadecane, isopropanol, toluene, petroleum ether, N, N-Dimethyl formamide, chloroform, acetone, isoamyl alcohol and isobutanol) at 10% (v/v) under standard assay conditions. Water was used as the solvent in the control reaction. Substrate specificity of the purified protease was determined using casein, bovine serum albumin and gelatin as substrates at 1% concentration under standard assay conditions.

The condition having the maximum activity of protease was considered as 100% for each experiment, and the activities in other conditions were plotted as relative percentages.

### 2.7 Enzyme kinetics

The rate of enzymatic reaction was measured by using casein as a substrate. The purified enzyme was incubated with different concentrations of buffered casein ranging from 0.1% to 1% for 30 minutes at 37 °C and protease activity was assayed using above mentioned protocol (Section 2.3). A Lineweaver-Burk plot showing 1/V_o_ (Y-axis) vs 1/[S] (X-axis) was plotted to calculate Michaelis-Menten constant (Km) and maximum reaction velocity (Vmax) values.

### 2.8 Protease identification by mass spectrometry

The purified protease was separated on 12% SDS-PAGE and visualized using silver staining. The protein band was excised from gel and digested with trypsin using trypsin profile IGD kit (Sigma-Aldrich, India). The peptides obtained after trypsin digestion were analyzed by Matrix-assisted Laser Desorption Ionization-Time of Flight (MALDI-TOF) analyzer (Applied Biosystems, USA). The tandem mass spectrometry (TOF/TOF) data obtained was evaluated by online software search engine i.e. MASCOT by searching in Bacteria (Eubacteria) database. The sequence of the identified protein was retrieved from UniProt (UniProt Consortium, 2014) and used for domain and sequence analysis. The domain analysis was done using Conserved Domain Database (Marchler-Bauer et al., 2014), Pfam (Bateman et al., 2004) and Interpro (Hunter et al., 2008). Its structure was predicted using homology modeling by Swiss model using structure of secreted protease C (1K7Q) as a template having more than 30% identity (Schwede et al., 2003).

### 2.9 Biofilm removal

#### 2.9.1 Biofilm formation

##### Staphylococcus aureus

MTCC 11949 procured from Microbial Type Culture Collection, Pune, India was grown in Luria-Bertani (LB) broth for biofilm formation assays. To study maximum biofilm formation, various media such as LB broth, nutrient broth, tryptone water and modified basal media (0.5% glucose, 0.5% peptone and 0.5% sodium chloride) were used for biofilm assays. The duration of biofilm formation was optimized by incubating *S. aureus* in various media for 24, 48 and 72 hours at 37 °C in 12 well tissue culture (TC) plates (1 ml per well) under stationary conditions to observe the maximum formation of biofilm.

#### 2.9.2 Biofilm staining

The effect of protease on bacterial biofilm was investigated by inoculating 1 ml of *S. aureus* culture (0.1 OD_600_) grown in LB medium in 12 well plate (Corning Inc., India). The plate was incubated at 37 °C for 72 hours to allow formation of biofilm. After 72 hours, the LB medium in each well of plate was replaced with fresh LB medium containing protease (10 μg/ml). This plate was further incubated for 24 hours where a well containing only LB medium without protease was considered as control. Congo red solution (final concentration of 50 μg/ml) was used to stain the preformed biofilm (Baidamshina et al., 2017).

The quantification of biofilm formation was done by using modified crystal violet assay (Sharma et al., 2015). 100 μl of bacterial suspension (0.1 OD_600_) was inoculated in 96-well polystyrene microtiter plate (Tarsons Product Pvt. Ltd., India) for 72 hours to allow biofilm formation. The used medium was replaced with fresh LB medium containing protease at different concentrations (10 μg/ml, 100 μg/ml and 1000 μg/ml) followed by incubation for 24 hours at 37 °C. For crystal violet staining, culture supernatant was removed and the wells were washed gently with phosphate buffered saline (PBS) to remove loosely bound bacterial cells. The adherent cells were fixed by using 100 μl of methanol (HiMedia, India) for 20 minutes after which the plate was air dried. The fixed biofilms were stained with 100 μl of 1% crystal violet (HiMedia) in distilled water for 20 minutes. The plate was washed with water to remove the excess stain. The cell bound crystal violet was extracted using 100 μl of 30% glacial acetic acid (HiMedia, India) in distilled water, and the absorbance measured at 595 nm using a Multiskan Ascent microplate reader (Thermo, Electron Corporation, USA). Cell free medium was used as a control.

#### 2.9.3 Assessment of antibacterial activity

Minimum inhibitory concentration (MIC) and minimum bactericidal concentration (MBC) of antibiotic (kanamycin) on *S. aureus* MTCC 11949 was evaluated using standard protocols (Baidamshina et al., 2017). MIC was evaluated as the lowest concentration of kanamycin that was sufficient to prevent visible bacterial growth (no turbidity) after 24 hours of incubation. MIC of kanamycin was evaluated by broth microdilution method to vary the concentration of kanamycin (256-0.5 μg/ml) by two fold serial dilution. These wells were inoculated with 200 μl of bacterial suspension (0.1 OD_600_) followed by incubation at 37 °C for 24 hours. The plates were then checked for visible bacterial growth.

MBC was determined as the lowest concentration of kanamycin necessary to kill 99.9% of bacteria. This is based on subculturing of bacterial inoculum (5 μl) from the MIC wells that showed no visible growth into 5 ml of fresh LB broth followed by incubation for 24 hours at 37 °C (Baidamshina et al., 2017) in a shaker incubator.

#### 2.9.4 Accessibility of antibiotic in biofilm

The accessibility of antibiotic in biofilm matrix in the presence and absence of protease was evaluated using drop plate assay (Herigstad et al., 2001) and confocal microscopy (Peak et al., 2010).

##### 2.9.4.1 Drop plate assay

The biofilm was formed by incubating staphylococcal culture for 72 hours as described above (Section 2.9.1). The biofilms were treated with either protease (10 μg/ml), or antibiotic (at 1x, 2x, 4x and 8x MBC concentrations) or a combination of protease and antibiotic for 24 hours at 37 °C. The wells were washed gently with 0.9% sodium chloride for removal of non-adherent cells. The cells entrenched in biofilms were suspended in 0.9% sodium chloride by scraping the bottom of the wells by repeated pipetting to help breakdown of bacterial aggregates. 100 μl of planktonic bacterial suspension was pipetted into an eppendorf having 900 μl of sterile 0.9% sodium chloride. 50 μl of this suspension was dispensed on LB agar plates as 10 μl drops. The last two drops were used to enumerate the colony forming units having countable number of colonies.

#### 2.9.4.2 Biofilm assay with confocal microscopy

The viability of biofilm-embedded cells was evaluated by modified live/ dead staining method using confocal microscopy (Peak et al., 2010). In this assay, biofilms were formed on coverslips in 12 well plates upon 72 hours incubation. After biofilm formation, cells were treated in a similar set of experiments as mentioned in drop plate assay except that only one concentration of antibiotic (8x MBC) was used in confocal microscopy.

After treatment for 24 hours at 37 °C, the plates were decanted and the glass coverslips were washed with phosphate buffered saline (PBS) to remove non-adherent cells. The samples were stained for 15 minutes with fluorescein diacetate (Sigma-Aldrich, India) and propidium iodide (Sigma-Aldrich, India) each at a final concentration of 10 μg/ml. After staining, the cells were washed twice with phosphate buffered saline (PBS) and mounted on glass slides using fluoromount (Sigma-Aldrich, India). Images were captured using Nikon A1R confocal microscope at 60x magnification.

#### 2.9.5 Cytotoxicity assay

Human epidermoid carcinoma cells (A431 cells) were obtained from National Centre for Cell Science, Pune, India. The cells were cultured in Dulbecco’s Modified Eagle Medium supplemented with 10% fetal bovine serum and 1X antibiotic-antimycotic mix (Gibco Thermo Fisher Scientific, India) at 37 °C and 5% CO_2_. The cells were seeded (20,000 cells/ml) in 96-well tissue culture plates and incubated overnight. Cell viability was tested in the presence of protease at various concentrations using 3-(4,5-dimethylthiazol-2-yl)-2,5-diphenyltetrazolium bromide (MTT) assay (Hansen et al., 1989). MTT (yellow) is bioreduced by mitochondrial dehydrogenase of viable cells into a purple colored product (formazan) which was measured at 595 nm.

The morphological changes in A431 cells due to treatment with purified protease were studied by phase contrast microscopy using Evos FL cell imaging system. The effect of protease treatment on A431 cells was analyzed for 24 and 48 hours by growing them on glass coverslips in the presence of protease. Treated cells were stained and visualized as above (Section 2.9.4.2).

### 2.10 Statistical analysis

Values in figures 3, 5 and 7 are represented as mean□±□standard error of mean acquired from three replicates.

**Figure 1.**
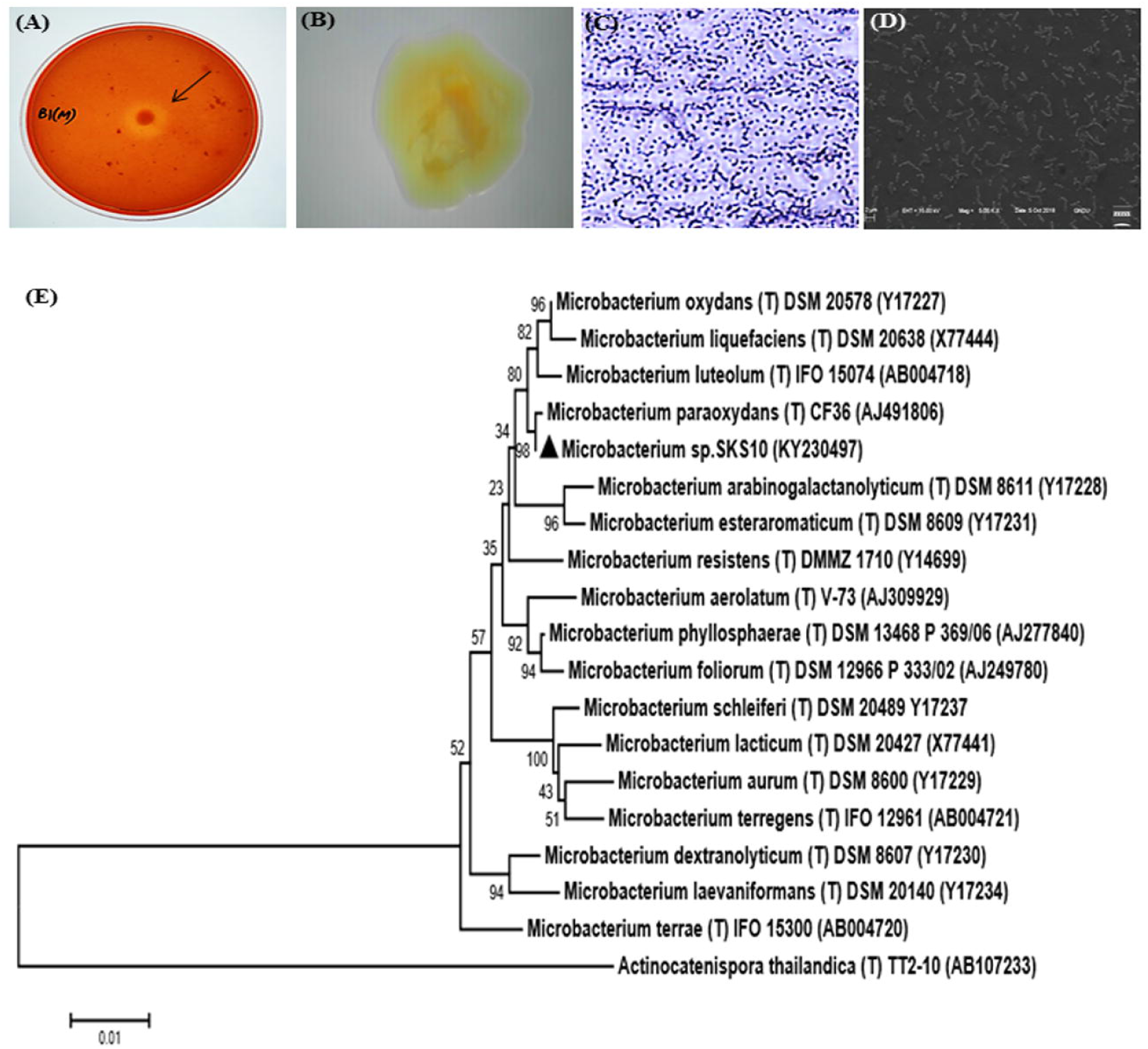
Morphological characterization of *Microbacterium* sp. SKS10. (A) Clear zone surrounding a colony of *Microbacterium* sp. SKS10 showing protease activity on skim milk agar. (B) Colony of *Microbacterium* sp. SKS10 as seen by a stereomicroscope. (C) Gram staining of *Microbacterium* sp. SKS10 showing gram-positive rods. (D) SKS10 observed by scanning electron microscope (10, 000x). (E) A phylogenetic tree of *Microbacterium sp*. SKS10 and its closest homologs based on their 16S rRNA sequences using Mega 7 software by using neighbor-joining method having branch length of 0.23688825. The percentage of bootstrap values based on 500 replicates is indicated at the branch nodes.

**Figure 2.**
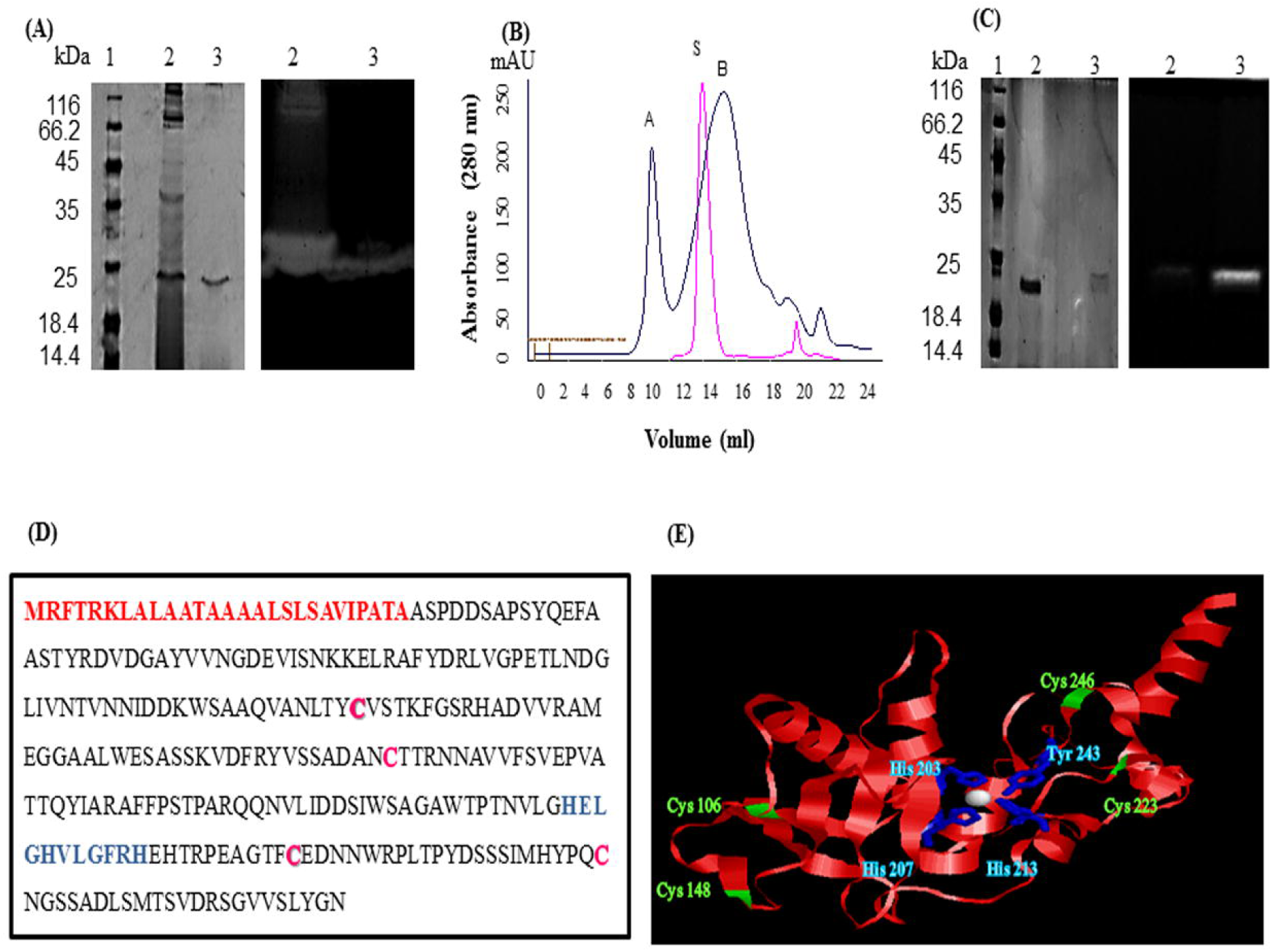
Purification and identification of protease from *Microbacterium* sp. SKS10. (A) 12% SDS-PAGE (left) and casein zymogram (right) of ammonium sulphate precipitated and GPC purified protease [Lane 1: molecular weight marker, lane 2: ammonium sulphate precipitated fraction, lane 3: GPC purified protease (peak B from figure 2B)]. (B) Overlay of elution profiles of protease (black) and chymotrypsin (red) on gel permeation chromatography. (C) 12% SDS-PAGE (left) and casein zymogram (right) of protease under reducing and non-reducing conditions. (Lane 1: molecular weight marker, lane 2: protease in reducing conditions, lane 3: protease in non-reducing conditions). (D) Protein sequence of protease identified as Peptidase M16 by mass spectrometry. Signal peptide (1-27) is shown in red, cysteine residues are in pink whereas HEXXHXXGXXH motif is highlighted in blue. (E) Structure modeled by Swiss-model using secreted protease C as a template. Cysteine residues are shown in green and residues His 203, His 207, His 213 and Tyr 243 chelating Zn (white sphere) are shown in blue.

**Figure 3.**
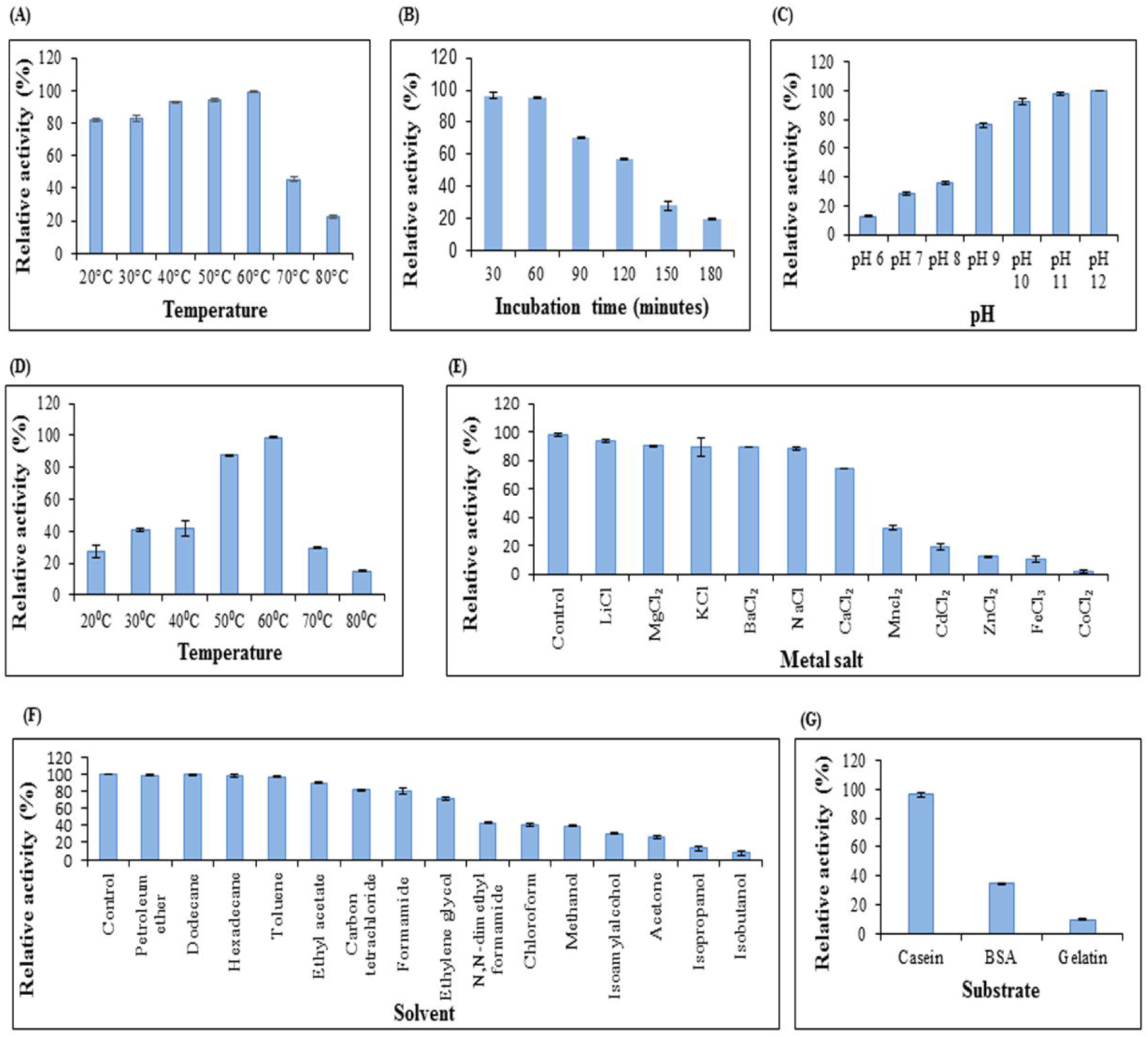
Characterization of protease. (A) Temperature stability of protease. Enzyme was preincubated at various temperatures followed by modified Lowry assay at 37 °C. (B) Enzyme activity at 37 °C after preincubation of protease at 60 °C to determine stability of protease at 60 °C. (C) Effect of pH on protease activity. Assay was performed at different pH values at 37 °C. (D) Effect of temperature on enzyme activity. Assay was performed at pH 12 with varying temperatures. (E) Effect of chloride metal salts (10 mM) on enzyme activity at pH 12, 37 °C. (F) Effect of solvents (10% v/v) on enzyme activity at pH 12, 37 °C. (G) Protease activity on different substrates. Columns and error bars above column represent mean ± standard error of the means respectively.

**Figure 4.**
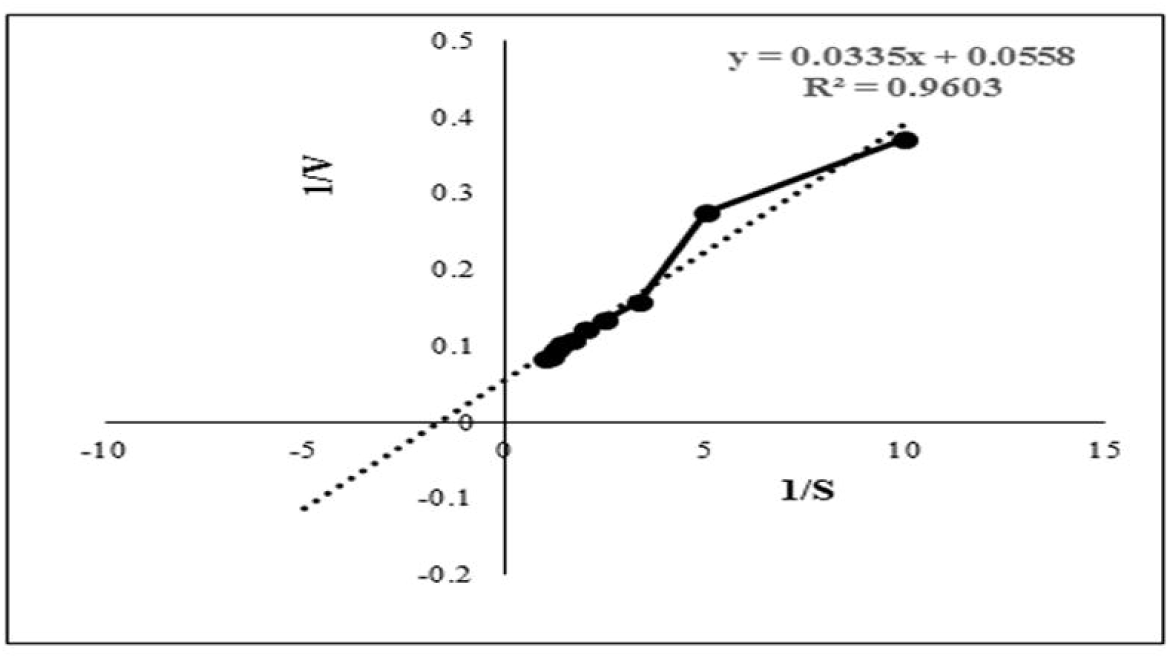
Lineweaver-Burk plot used to determine the kinetic parameters such as Km and Vmax. Intercepts on X and Y axis were used to determine 1/Km and 1/Vmax respectively.

**Figure 5.**
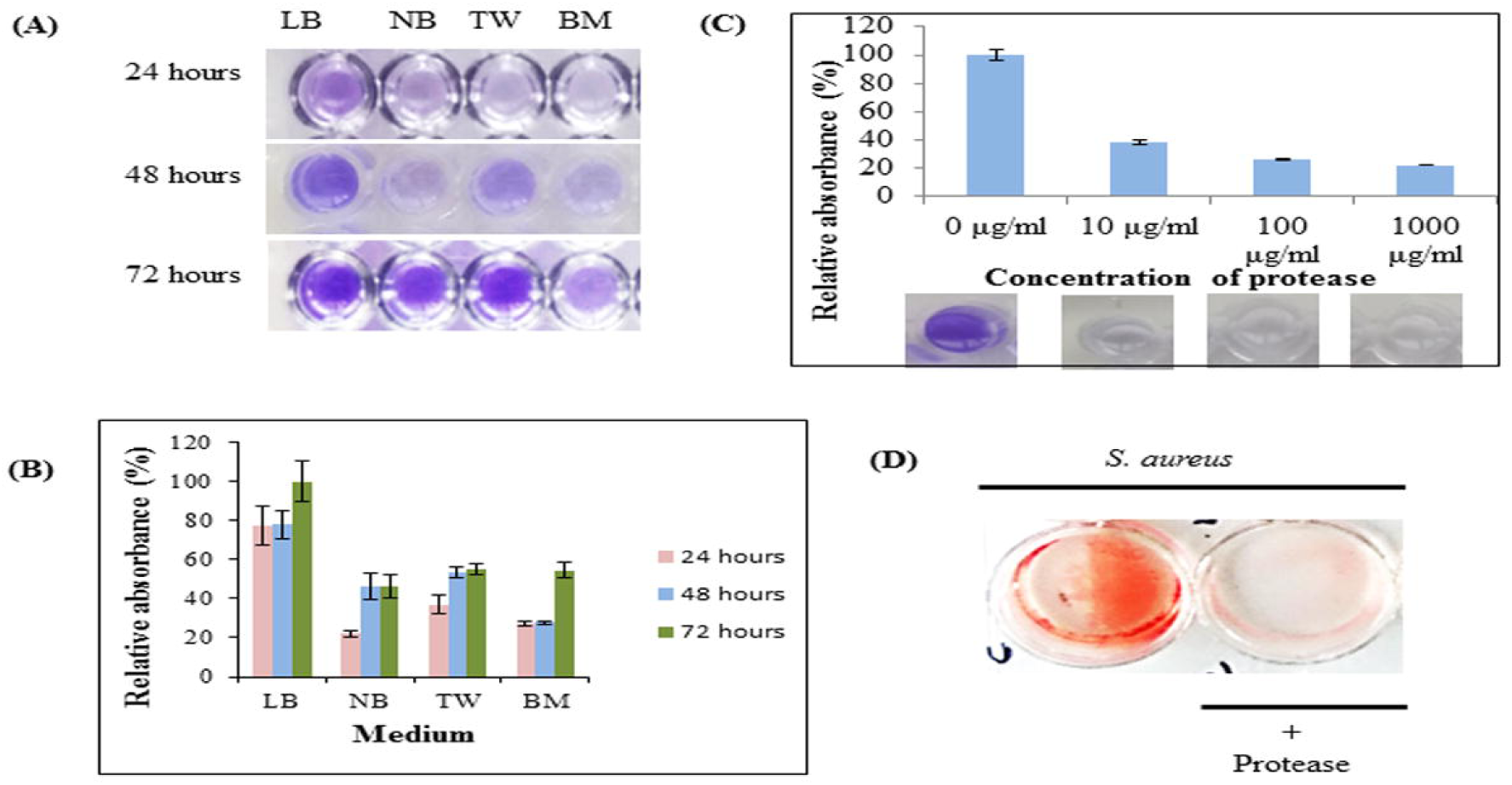
Protease as an anti-biofilm agent. (A) The biofilm formation by *S. aureus* cultivated in Luria Bertani broth (LB), nutrient broth (NB), tryptone water (TW) and basal medium (BM) on 35-mm microtiter plates for 24, 48 and 72 hours as determined by crystal violet staining. (B) Relative absorbance (595 nm) from crystal violet stained biofilms solubilized with 30% glacial acetic acid. Experiment was performed in triplicates and an average of relative absorbances (%) plotted. Maximum absorbance obtained in the experiment was considered as 100%. (C) Dispersal of staphylococcal biofilm on treatment with protease at final concentrations of 10 μg/ml, 100 μg/ml and 1000 μg/ml for 24 hours as quantified by crystal-violet staining. Top panel shows an average of relative absorbances (595 nm) as calculated in figure 5B. Bottom panel shows a representative well each from these experiments. (D) Dispersal of preformed biofilm by protease stained by congo red. An untreated well (without protease) incubated under identical conditions and stained with congo red was considered as control (red).

**Figure 6.**
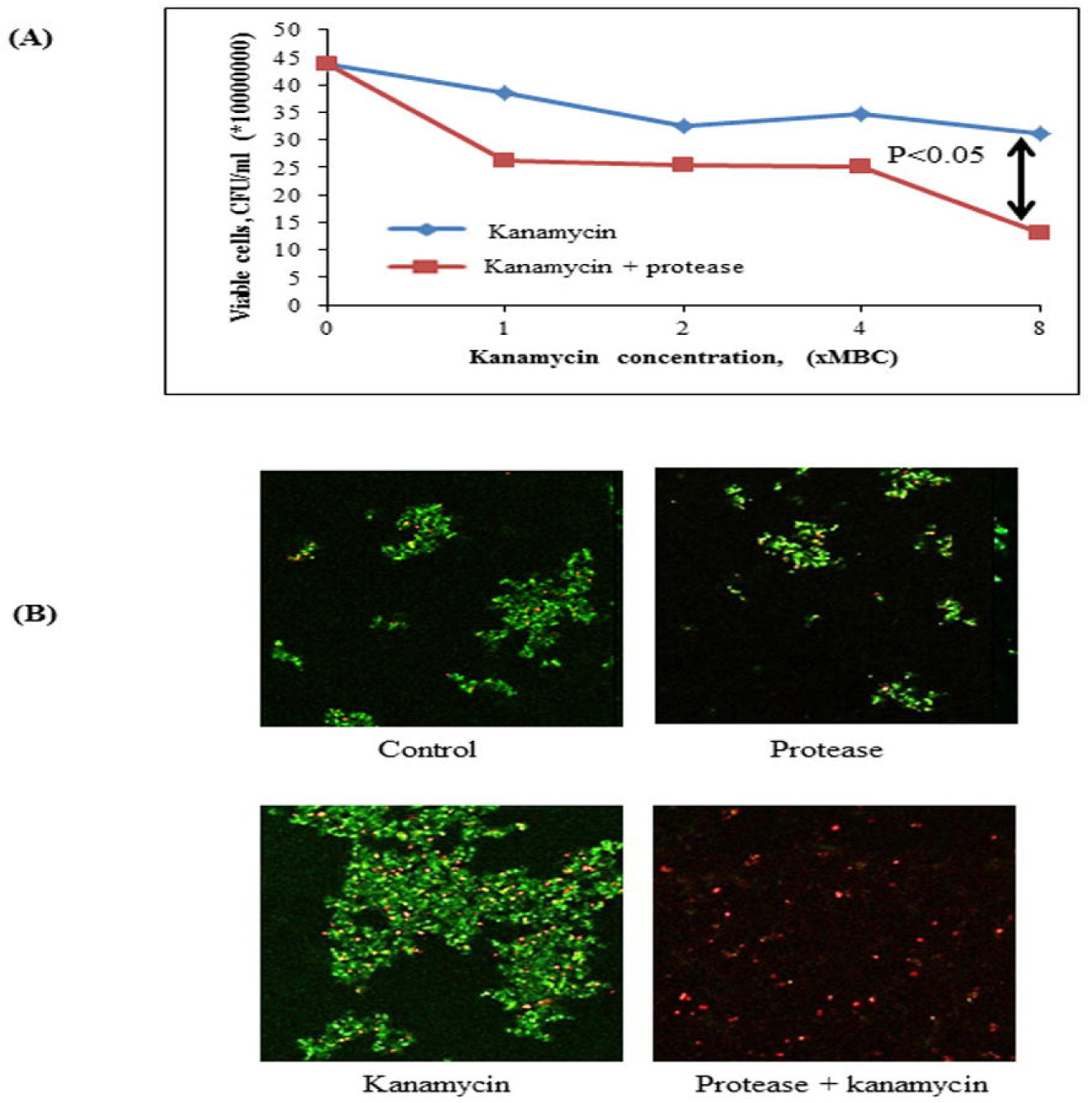
Effect of protease treatment on accessibility of kanamycin against biofilm embedded Staphylococci. (A) Cell viability post treatment (24 hours) with kanamycin (8x MBC) or kanamycin (8x MBC) + protease (10 μg/ml) in a drop plate assay. The adherent cells were scraped and suspended in sterile 0.9% NaCl, dispensed as 10 µl drops on LB agar and CFUs counted from last two drops. (B) Preformed biofilms of *S. aureus* on glass coverslips were incubated for 24 hours in the presence of protease (10 μg/ml) and kanamycin (8x MBC), and analyzed with confocal microscopy after live (green)/ dead (red) staining using fluorescein diacetate and propidium iodide.

**Figure 7.**
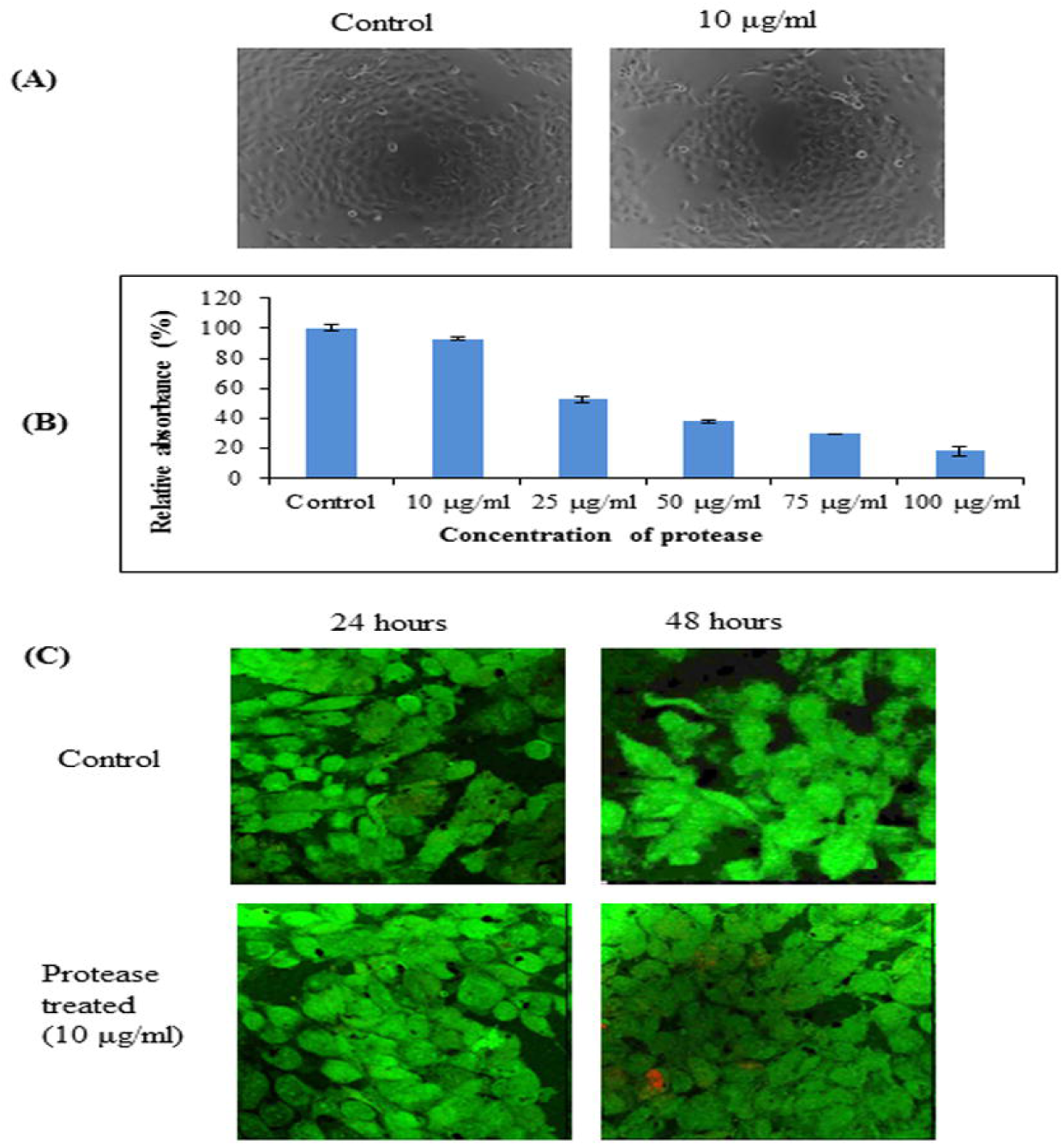
Cytotoxicity of protease on A431 cells. (A) Representative phase contrast photo micrographs of A431 cells in control and cells treated with protease (10 µg/ml) for 24 hours. (B) Bar diagrams representing cytotoxicity by MTT assay on A431 cells treated with 10-100 µg/ml of protease. (C) Proliferation of human epidermoid cells analyzed by live (green)/ dead (red) staining after 24 and 48 hours of protease treatment studied by confocal microscopy.

## 3 Results and discussion

### 3.1 Isolation, identification and characterization of *Microbacterium* sp. SKS10

An alkaline protease producing bacterial strain was isolated from soil of west coast region of India. Colonies of bacteria showed a zone of clearance on skim milk agar plates (pH 10) stained with congo red (Figure 1A). The shape and morphology of bacterial colonies was observed as entire, opaque, smooth and yellowish-orange on nutrient clerigel plates under a stereomicroscope (Figure 1B). The cells were gram-positive and short rods as observed by gram staining (Figure 1C) and scanning electron microscope (Figure 1D) respectively. The molecular characterization of the strain was performed by sequencing of PCR amplified 16S rRNA gene using universal bacterial primers. The sequence of 16S rRNA having 1331 base pairs was deposited to GenBank with accession number KY230497 (https://www.ncbi.nlm.nih.gov/nuccore/KY230497.1). The sequence was searched in RDP-II, EzTaxon and GenBank database using BLAST. BLASTn result at National Centre for Biotechnology Information (NCBI) database showed 100% sequence identity to a sequence of *Microbacterium paraoxydans* strain DAS46. Results from EzTaxon, a database for 16S ribosomal DNA sequences of “type strains” showed 99.77% sequence identity to *Microbacterium paraoxydans* NBRC 103076 (T). Similarity search against RDP-II, a database of 16S ribosomal DNA sequences of both “type and non-type strains” showed 99.45% identity to *Microbacterium paraoxydans* CF36 (T). This strain belongs to the phylum Actinobacteria, class Actinobacteria, subclass Actinobacteridae, order Actinomycetales, suborder Micrococcineae, family Microbacteriaceae and genus Microbacterium. The phylogenetic analysis using sequences of nearest homologs retrieved from RDPII database showed presence of *Microbacterium sp.* SKS10 in a cluster with *Microbacterium paraoxydans CF36* (T) in the phylogenetic tree constructed using neighbor-joining method (Figure 1E).

The biochemical characterization of strain SKS10 was performed using HiCarbo kit. *Microbacterium* sp. SKS10 was able to utilize glycerol, salicin, esculin, citrate and malonate among all the carbohydrates available in HiCarbo kit (Table S1). Laffineur et al. (2003) reported that *Microbacterium paraoxydans* CF36, the nearest homolog of SKS10 in RDPII database was able to utilize lactose, maltose, fructose, mannose, □-methyl-D-glucoside, Rhamnose and D-arabinose whereas *Microbacterium* sp. SKS10 was unable to utilize these indicating the difference between two strains. The strain SKS10 could produce an extracellular protease but was unable to secrete other enzymes such as amylase, lipase and lysine decarboxylase (Table S2). This strain did not secrete stable acids, acetylmethylcarbinol and indole as observed by methyl red test, vogues proskauer and tryptone water test respectively whereas no information regarding these tests was provided by Laffineur et al. about the strain CF36 (Table S2).

### 3.2 Protease purification

The crude enzyme precipitated from the culture supernatant of 70% ammonium sulphate fraction resulted in a yield of 65.717% and specific activity of 34.847 U/mg (1.13 fold purification) (Table 1). A zymogram of this preparation showed two clear zones corresponding to a high (>66 kDa) and a low molecular weight (∼25 kDa) suggestive of the presence of two proteases (Figure 2A). The partially purified proteins obtained after ammonium sulphate precipitation were further purified by gel permeation chromatography (GPC) where chymotrypsin (25 kDa) was used as a standard. Two peaks eluted before (A) and after (B) chymotrypsin (Figure 2B). Peak A showed multiple bands on SDS-PAGE (data not shown) and was not used for further analysis. Peak B showed a single band (∼25 kDa) on silver stained SDS-PAGE and had protease activity as depicted by a single clear zone in the zymogram (Figure 2A). The specific activity and yield of the protease in this fraction were 160.21 U/mg and 1.77% respectively with 5.21 fold purification. This fraction was stored at - 20 °C and used for all enzyme characterization and biofilm assays.

**Table 1.** Purification of protease from *Microbacterium* sp. SKS10.

| **Purification step** | **Total activity (U)<sup>a</sup>** | **Total protein (mg)<sup>b</sup>** | **Specific activity (U/mg)** | **Yield (%)** | **Fold purification** |
| --- | --- | --- | --- | --- | --- |
| Crude enzyme | 2004.8 | 65.2 | 30.748 | 100 | 1 |
| Ammonium sulphate precipitation | 1317.5 | 37.809 | 34.847 | 65.7 | 1.13 |
| Gel permeation chromatography | 35.568 | 0.222 | 160.21 | 1.77 | 5.21 |

<sup>a</sup>Enzyme activities were determined by using casein as a substrate spectrophotometrically at 660 nm
<sup>b</sup>The protein concentration was estimated using Bradford's method

A difference in mobility of the purified enzyme on SDS-PAGE under reducing and non-reducing conditions was observed indicating the presence of intra-chain disulphide bonds. The protease activity was lost under reducing conditions as visible in the zymogram proving that these disulphide linkages are important for protease activity (Figure 2C).

### 3.3 Identification of protease

A single protein band from silver stained SDS-PAGE was excised, trypsinized and used for mass spectrometry to identify the protease. The mass fingerprint obtained by tandem mass spectrometry was searched in bacterial protein database using MASCOT (Figure S1) which resulted in Peptidase M16 (*Microbacterium*) as the first hit (Figure 2D). The amino acid sequence of this peptidase (UniProt ID: A0A1H1U687) (molecular weight 29 kDa) was retrieved from the database. Removal of the signal sequence (1-27) present at the N-terminus (Figure 2D) of this protein would result in a processed protease (28-269) carrying a molecular weight similar to that observed in SDS-PAGE and GPC (Figures 2A,B). Domains and motifs predicted using various databases (Table 2) revealed the presence of a zinc-binding motif i.e. HEXXHXXGXXH (Figure 2D) in its sequence. Four cysteine residues in the protein sequence seemed to be involved in disulphide bond formation (C 106, C246, C148, C223). The presence of disulphide bonds is a characteristic of Astacin family of metallopeptidases, and the HEXXH motif and metallopeptidase activity is found in Astacin, ZnMc, Peptidase_M57, IPR024653 and IPR006026. The structure of protein modeled using secreted protease C as a template (Figure 2E) suggested that residues such as His 203, His 207, His 213 and Tyr 243 bind to a zinc ion (Figure 2E). Cys 106 and Cys 246 are present in close proximity to Cys 148 and Cys 223 respectively, suggesting that these pairs may form disulphide bonds. Our results suggest this protease to be a metallo-peptidase.

**Table 2.** List of domain hits with their description.

| **Database** | **Domain hits**<br>**(Interval)** | **Description** |
| --- | --- | --- |
| **Conserved**<br>**Domain**<br>**database- NCBI**<br>(Marchler-Bauer<br>et al., 2014) | ZnMc_BMP1_TLD<br>(203-243) | Zinc dependent metalloproteases |
|  | Astacin (203-243) | Requires zinc for catalysis and has two conserved disulphide bridges. It has N-terminal propeptide which cleaves to release active protease. |
|  | ZnMc (94-218) | Zinc dependent metalloprotease having HEXXH zinc binding site |
|  | Peptidase M_10<br>(102-268) | Cleaves peptides and require zinc for catalysis |
| **Pfam** (Bateman<br>et al., 2004) | Peptidase_M57<br>(85-267) | Dual action HEIGH metallopeptidase |
| **InterPro** (Hunter<br>et al., 2008) | IPR024079<br>(77-269) | Metallopeptidase activity |
|  | IPR024653<br>(114-214) | Represents metallopeptidase M10, M27 and M57. The catalytic triad is HE-H-H and many members have the sequence motif HEIGH. |
|  | IPR006026<br>(91-246) | Presence of HEXXH motif which forms a part of metal binding site |

### 3.4 Characterization of protease

The GPC purified metalloprotease having specific activity of 160.21 U/mg was used for protease characterization (Figure 3). The protease was observed to be stable at a wide range of temperatures ranging from 20-60 °C on pre-incubation of the enzyme at temperatures ranging from 20 °C-80 °C (Figure 3A). The protease was determined to be stable at 60 °C as it retained 95% and 57% of its activity after incubation at this temperature for one and two hours respectively (Figure 3B). Prolonged incubation (180 minutes) at 60 °C led to 80% decline in enzyme activity. The protease showed its maximum activity at pH 12 and 60 °C when the reaction mixture was incubated at pH varying from 6 to 12 (Figure 3C) and a range of temperatures varying from 20 °C to 80 °C (Figure 3D) respectively. These results suggest its application in industries requiring moderate heat and alkaline conditions.

This protease was quite stable in chloride salts of lithium, magnesium, potassium, barium and sodium as it retained more than 85% of its activity in their presence whereas the enzyme activity decreased to 1.88% in the presence of cobalt (II) chloride (Figure 3E). The protease also retained more than 85% activity in the presence of petroleum ether, dodecane, hexadecane, toluene and ethyl acetate using solvents at 10% concentration (v/v) whereas activity got declined to 13.31% and 7.41% in the presence of isopropanol and isobutanol respectively (Figure 3F). In a similar report, the keratinolytic serine protease produced by *Streptomyces* sp. strain AB1 showed stability in the presence of ethyl acetate whereas isopropanol and butanol inhibited the enzyme activity (Jaouadi et al., 2010).

The effect of enzyme action on various substrates was analyzed. It was observed that the protease showed its maximum activity (100%) when casein was used as a substrate (Figure 3G) and retained up to 34.22% and 9.678% activity using bovine serum albumin and gelatin respectively. The proteases secreted by *Chryseobacterium indologenes* TKU014 also showed higher activity towards casein (Wang et al., 2008).

The Km and Vmax values of this protease from *Microbacterium* sp. SKS10 were calculated as 0.577 mg/ml and 17.2413 U/ml respectively using casein as a substrate by Lineweaver-Burk plot (Figure 4). Km of a protease secreted by *Rhizopus oryzae* (7 mg/ml) was high (Mushtaq et al., 2015) proving that the protease used in the present study has a higher substrate affinity as compared to the protease reported by Mushtaq et al. However, Km values were quite comparable with that of a thermoactive protease secreted by *Bacillus* sp. (Jain et al., 2012).

### 3.5 Biofilm removal by protease

Various chemical strategies have been in use for biofilm removal since years but these chemicals possess various health as well as environmental hazards. Due to limitations of chemicals used for biofilm removal, chemical strategies can be replaced with enzymatic methods.

In this study, the protease was able to degrade biofilms formed by *S. aureus* as observed by modified crystal violet assay (Figure 5). The formation of staphylococcal biofilm was studied in various media such as LB broth, nutrient broth, tryptone water and basal media for 24, 48 and 72 hours (Figure 5A). It was observed that staphylococcal biofilm was best formed in LB medium (Figure 5B). This protease was able to degrade biofilms up to 61.923%, 73.732% and 77.728% on treatment at 10 μg/ml, 100 μg/ml and 1000 μg/ml concentrations respectively (Figure 5C). Previous reports suggest that enzymes (at concentration of 1 mg/ml) such as papain, trypsin (Trizna et al., 2016), □-Amylase (Craigen et al., 2011) and DNaseI (Tetz et al., 2009) were able to degrade biofilms by 50-60% (Table 3). Our results showed that this metalloprotease was effective in biofilm degradation at 100 times lower concentration as compared to these enzymes.

**Table 3.** Enzymatic removal of biofilms.

| S. No. | Enzyme | Concentration | % Biofilm removal | References |
| --- | --- | --- | --- | --- |
| 1 | Papain or Trypsin | 100 µg/ml<br>1000 µg/ml | 20-30%<br>50-60% | Trizna et al., 2016 |
| 2 | □-Amylase | 1 mg/ml<br>2 mg/ml<br>10 mg/ml<br>20 mg/ml<br>100 mg/ml | 64%<br>68%<br>80%<br>81%<br>82% | Craigien et al., 2011 |
| 3 | rChymotrypsin | 10 µg/ml | 51% | Harris et al., 2013 |
| 4 | DNaseI | 1 mg/ml | 70% | Tetz et al., 2009 |
| 5 | AcylaseI | 5 µg/ml | 9% | Kim et al., 2013 |
| 6 | Proteinase K | 100 µg/ml | 56.6% | Kim et al., 2013 |
| **7** | **Protease** | **10 µg/ml** | **62%** |  |

The *S. aureus* biofilm matrix consists of host factors, secreted and lysis-derived proteins, polysaccharides and DNA. Hydrolysis of protein components from the extracellular matrix of biofilms by protease was observed using congo red, a dye used to stain amyloid proteins (Figure 5D). The control wells (only bacteria in absence of protease) stained red with congo red whereas presence of protease showed a decrease in staining intensity signifying degradation of the extracellular matrix.

#### 3.5.1 Enhancement of accessibility of antimicrobials by protease

The bacteria embedded into matrix of biofilm are inaccessible to antibiotics. The enhancement of accessibility of antimicrobials by protease treatment was determined by drop plate assay and confocal microscopy. *S. aureus* is reported to be sensitive to kanamycin (Pengov and Ceru, 2003). MIC and MBC values of kanamycin were determined by broth microdilution method for planktonic *S. aureus* MTCC 11949 as 4 μg/ml and 8 μg/ml respectively. Kanamycin, when used alone at various concentrations (1x MBC-8x MBC) did not affect the viability of bacterial cells in biofilm probably due to poor penetration of antibiotic in the biofilm (Figure 6A). The viability of bacterial cells in the biofilm decreased significantly on treatment of cells with kanamycin (8x MBC) in the presence of protease (10 μg/ml). This may be due to better penetration of kanamycin in biofilm after degradation of the protein backbone of extracellular matrix by protease treatment, thus killing the bacteria embedded in biofilms (Table 4). Treatment of biofilm with protease (10 μg/ml) alone did not affect the viability of bacteria as tested in a drop plate assay, demonstrating that it was non-toxic to cells (Figure 6A). Cell viability was also observed by live/dead staining using fluorescein diacetate and propidium iodide by confocal microscopy. Cells treated with protease (10 μg/ml) and kanamycin (8x MBC) alone stained green suggesting their viability, while cells treated with same concentrations of kanamycin and protease in combination stained red depicting dead cells (Figure 6B).

**Table 4.**
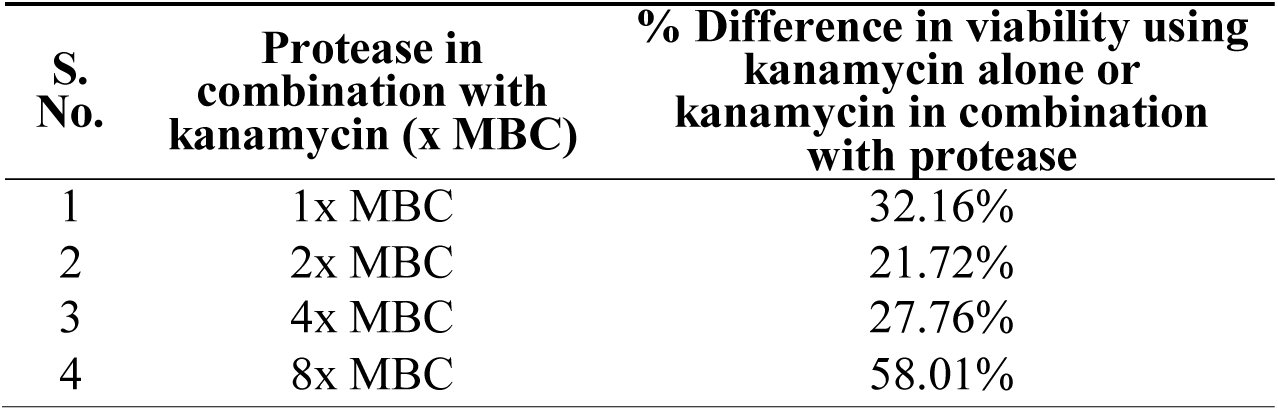
Percent difference in viability of cells in biofilm on treatment with kanamycin (× MBC) alone or kanamycin in combination with protease

#### 3.5.2 Cytotoxicity evaluation

Cytotoxicity of the protease was investigated on human epidermoid carcinoma A431 cells by phase contrast microscopy, MTT assay and confocal microscopy. It was observed that there was no change in morphological characteristics (Figure 7A) of A431 cells upon treatment with protease (10 μg/ml). Also, there was no significant reduction in dehydrogenase activity of cells treated with protease (92.76% retained) at a concentration of 10 μg/ml for 24 hours as seen by MTT assay (Figure 7B). However, higher concentrations (>10 μg/ml) were found to be toxic to human epidermoid cells (Figure 7B).

The human epidermoid carcinoma cells were live/ dead stained and analyzed by confocal microscopy after protease treatment for 48 hours (Figure 7C). There was no significant increase in necrotic cells in either control or protease containing wells at 10 μg/ml concentrations suggesting it to be safe for application on human cells at this concentration.

## 4 Conclusions

This study extrapolates the potential of an extracellular protease secreted by *Microbacterium* sp. SKS10. The protease was purified with ammonium sulphate precipitation followed by gel permeation chromatography. This protease (25 kDa) identified as Peptidase M16 using mass spectrometry showed its optimum activity at 60 °C and pH 12, and retained activity in various chloride metal salts as well as organic solvents. The loss of protease activity was observed under reducing conditions and also in the presence of cobalt chloride. The protease has a potential for biofilm removal when used at 100 times lower concentration as compared with other common enzymes. As per our knowledge, this is the first report of the use of Peptidase M16 as an anti-biofilm agent which increased the penetration of antibiotics by degrading the bacterial extracellular matrix. The enzyme was non-cytotoxic to human epidermoid cells suggesting its use in biofilm removal and in therapeutic and industrial applications in the future.

## Author Contributions

SKS and PCM were involved in the study design. SKS carried out the experiments. GJ was involved in mass spectrometry studies. SKS and PCM wrote the manuscript. All authors have read and approved the final manuscript.

## Abbreviations

NCBI: national center for biotechnology information
BLAST: basic local alignment search tool
RDPII: ribosomal database project II
PCR: polymerase chain reaction
SDS-PAGE: sodium dodecyl sulphate-polyacrylamide gel electrophoresis
GPC: gel permeation chromatography
LB: luria broth
MIC: minimum inhibitory concentration
MBC: minimum bactericidal concentration
MTT: 3-(4,5-dimethylthiazol-2-yl)-2,5-diphenyltetrazolium bromide

## Acknowledgments

SKS is a UGC-MANF-SRF (University Grants Commission-Maulana Azad National Fellowship-Senior Research Fellow). We would like to thank Dr. Rachna Hora, Assistant Professor, Department of Molecular Biology and Biochemistry, Guru Nanak Dev University, Amritsar, Punjab, India for reading the manuscript.

## References

Algburi, A., Comito, N., Kashtanov, D., Dicks, L. M., and Chikindas, M. L. (2017). Control of biofilm formation: antibiotics and beyond. Appl. Environ. Microbiol. 83, e02508–16. doi: 10.1128/AEM.02508-16

Araujo, P., Lemos, M., Mergulhao, F., Melo, L. and Simoes, M. (2011). Antimicrobial resistance to disinfectants in biofilms. Science against microbial pathogens: communicating current research and technological advances, 3, 826–834.

Baidamshina, D. R., Trizna, E. Y., Holyavka, M. G., Bogachev, M. I., Artyukhov, V. G., Akhatova, F. S., Rozhina, E. V., Fakhrullin, R. F., and Kayumov, A. R. (2017). Targeting microbial biofilms using Ficin, a nonspecific plant protease. Sci. Rep. 7, 46068. doi:10.1038/srep46068

Bateman, A., Coin, L., Durbin, R., Finn, R. D., Hollich, V., Griffiths-Jones, S., Khanna, A., Marshall, M., Moxon, S., Sonnhammer, E. L. and Studholme, D. J. (2004). The Pfam protein families database. Nucleic acids Res. 32, D138–D141. doi: 10.1093/nar/gkh121

Bester, D. J., Kupai, K., Csont, T., Szucs, G., Csonka, C., Esterhuyse, A. J., Ferdinandy, P., and Van Rooyen, J. (2010). Dietary red palm oil supplementation reduces myocardial infarct size in an isolated perfused rat heart model. Lipids Health Dis. 9, 64. doi: 10.1186/1476-511X-9-64

Boominadhan, U., Rajakumar, R., Sivakumaar, P. K. V., and Joe, M. M. (2009). Optimization of protease enzyme production using *Bacillus* sp. isolated from different wastes. Bot. Res. Int. 2, 83–87.

Bradford, M. M. (1976). A rapid and sensitive method for the quantitation of microgram quantities of protein utilizing the principle of protein-dye binding. Anal Biochem. 72: 248–54. doi: https://doi.org/10.1016/0003-2697(76)90527-3

Chun, J., Lee, J. H., Jung, Y., Kim, M., Kim, S., Kim, B. K., and Lim, Y. W. (2007). EzTaxon: a web-based tool for the identification of prokaryotes based on 16S ribosomal RNA gene sequences. Int. J. Syst. Evol. Microbiol. 57, 2259–2261. doi: 10.1099/ijs.0.64915-0

Craigen, B., Dashiff, A., and Kadouri, D. E. (2011). The use of commercially available alphaamylase compounds to inhibit and remove *Staphylococcus aureus* biofilms. Open Microbiol. J. 5, p. 21. doi: 10.2174/1874285801105010021

Fleming, D., and Rumbaugh, K. P. (2017). Approaches to dispersing medical biofilms. Microorganisms. 5, 15. doi: 10.3390/microorganisms5020015

Gjermansen, M., Nilsson, M., Yang, L., and Tolker-Nielsen, T. (2010). Characterization of starvation-induced dispersion In *Pseudomonas putida* biofilms: genetic elements and molecular mechanisms. Mol. Microbiol. 75, 815–826. doi: 10.1111/j.1365-2958.2009.06793.x

Hansen, M. B., Nielsen, S. E., and Berg, K. (1989). Re-examination and further development of a precise and rapid dye method for measuring cell growth/cell kill. J. Immunol. Methods. 119, 203–210. doi: 10.1016/0022-1759(89)90397-9

Harris, L. G., Nigam, Y., Sawyer, J., Mack, D., and Pritchard, D.I. (2013). Lucilia sericata chymotrypsin disrupts protein adhesin-mediated staphylococcal biofilm formation. Appl. Environ. Microbiol. 79, 1393–1395. doi: 10.1128/AEM.03689-12

Herigstad, B., Hamilton, M., and Heersink, J. (2001). How to optimize the drop plate method for enumerating bacteria. J. Microbiol. Methods. 44, 121–129. doi: 10.1016/S0167-7012(00)00241-4

Hunter, S., Apweiler, R., Attwood, T. K., Bairoch, A., Bateman, A., Binns, D., Bork, P., Das, U., Daugherty, L., Duquenne, L. and Finn, R. D. (2008). InterPro: the integrative protein signature database. Nucleic acids Res. 37, D211–D215. doi: 10.1093/nar/gkn785

Jain, D., Pancha, I., Mishra, S. K., Shrivastav, A., and Mishra, S. (2012). Purification and characterization of haloalkaline thermoactive, solvent stable and SDS-induced protease from *Bacillus* sp.: a potential additive for laundry detergents. Bioresource Technol. 115, 228–236. doi: 10.1016/j.biortech.2011.10.081

Jamal, M., Tasneem, U., Hussain, T., and Andleeb, S. (2015). Bacterial biofilm: its composition, formation and role in human infections. RRJMB. 4, 1–15.

Jaouadi, B., Abdelmalek, B., Fodil, D., Ferradji, F. Z., Rekik, H., Zarai, N., and Bejar, S. (2010). Purification and characterization of a thermostable keratinolytic serine alkaline proteinase from *Streptomyces* sp. strain AB1 with high stability in organic solvents. Bioresource Technol. 101, 8361–8369. doi: 10.1016/j.biortech.2010.05.066

Johnson, M., Zaretskaya, I., Raytselis, Y., Merezhuk, Y., McGinnis, S., and Madden, T. L. (2008). NCBI BLAST: a better web interface. Nucleic acids Res. 36, W5–W9. doi: 10.1093/nar/gkn201

Kim, L. H., Kim, S. J., Kim, C. M., Shin, M. S., Kook, S., and Kim, I. S. (2013). Effects of enzymatic treatment on the reduction of extracellular polymeric substances (EPS) from biofouled membranes. Desalin. Water Treat. 51, 6355–6361. doi: 10.1080/19443994.2013.780996

Kumar, S., Stecher, G., and Tamura, K. (2016). MEGA7: molecular evolutionary genetics analysis version 7.0 for bigger datasets. Mol. Biol. Evol. 33, 1870–1874. doi: 10.1093/molbev/msw054

Laffineur, K., Avesani, V., Cornu, G., Charlier, J., Janssens, M., Wauters, G., and Delmee, M. (2003). Bacteremia due to a novel Microbacterium species in a patient with Leukemia and description of *Microbacterium paraoxydans* sp. nov. J. Clin. Microbiol. 41, 2242–2246. doi: 10.1128/JCM.41.5.2242-2246.2003

Lister, J. L., Horswill, A. R. (2014). *Staphylococcus aureus* biofilms: recent developments in biofilm dispersal. Front Cell Infect Microbiol. 4:178.

Lopez, D., Vlamakis, H., and Kolter, R. (2010). Biofilms. Cold Spring Harb. Perspect Bio. a000398. doi: 10.1101/cshperspect.a000398

Lowry, O. H., Rosebrough, N. J., Farr, A. L., and Randall, R. J. (1951). Protein measurement with the Folin phenol reagent. J. Biol. Chem. 193, 265–275.

Maidak, B. L., Cole, J. R., Lilburn, T. G., Parker Jr, C. T., Saxman, P. R., Stredwick, J. M., Garrity, G. M., Li, B., Olsen., G. J., Pramanik., S., and Schmidt, T. M. (2000). The RDP (ribosomal database project) continues. Nucleic acids Res. 28, 173–174.

Marchler-Bauer, A., Derbyshire, M. K., Gonzales, N. R., Lu, S., Chitsaz, F., Geer, L. Y., Geer, R. C., He, J., Gwadz, M., Hurwitz, D. I. and Lanczycki, C. J. (2014). CDD: NCBI’s conserved domain database. Nucleic acids Res. 43, D222–D226. doi: 10.1093/nar/gku1221

Mushtaq, Z., Irfan, M., Nadeem, M., Naz, M., and Syed, Q. (2015). Kinetics study of extracellular detergent stable alkaline protease from *Rhizopus oryzae*. Braz. Arch. Biol. Technol. 58, 175–184. doi: 10.1590/S1516-8913201400071

Pammi, N., Chaitanya, K., and Mahmood, S. K. 2015. Serratia marcescens OU50T sp. nov., a cellulose and pha producing bacterium isolated from polluted water. Int. J. Environ. Biol. 5: 32–36.

Peak, E., Chalmers, I. W., and Hoffmann, K. F. (2010). Development and validation of a quantitative, high-throughput, fluorescent-based bioassay to detect schistosoma viability. PLoS Negl. Trop. Dis. 4, e759. doi: 10.1371/journal.pntd.0000759

Pengov, A., and Ceru, S. (2003). Antimicrobial drug susceptibility of *Staphylococcus aureus* strains isolated from bovine and ovine mammary glands. J. Dairy Sci. 86, 3157–3163. doi: 10.3168/jds.S0022-0302(03)73917-4

Sambrook, J., Russell, D. W. and Russell, D.W. (2001). Molecular cloning: a laboratory manual (3-volume set) (Vol. 999). New York: Cold spring harbor laboratory press.

Schwede, T., Kopp, J., Guex, N. and Peitsch, M. C. (2003). SWISS-MODEL: an automated protein homology-modeling server. Nucleic acids Res. 31, 3381–3385. https://doi.org/10.1093/nar/gkg520

Sharma, D., Saharan, B. S., Chauhan, N., Procha, S., and Lal, S. (2015). Isolation and functional characterization of novel biosurfactant produced by *Enterococcus faecium*. SpringerPlus. 4, 4. doi: 10.1186/2193-1801-4-4

Singh, S., Singh, S. K., Chowdhury, I., and Singh, R. (2017). Understanding the mechanism of bacterial biofilms resistance to antimicrobial agents. Open Microbiol. J. 11, 53. doi: 10.2174/1874285801711010053

Sivanandhini, T., Subbaiya, R., Gopinath, M., Angrasan, J. K.V., Kabilan, T. and Selvam, M. M., 2015. An investigation on morphological characterization of Actinomycetes isolated from marine sediments. RJPBCS, 6(2), pp. 1234–1243.

Teitzel, G. M., Parsek, M. R. (2003). Heavy metal resistance of biofilm and planktonic *Pseudomonas aeruginosa*. Appl. Environ. Microbiol. 69, 2313–2320. doi: 0.1128/AEM.69.4.2313-2320.2003

Tetz, G. V., Artemenko, N. K., and Tetz, V.V. (2009). Effect of DNase and antibiotics on biofilm characteristics. Antimicrob. Agents Chemother. 53, 1204–1209. doi: 10.1128/AAC.00471-08

Trizna, E., Diana, B., Kholyavka, M., Sharafutdinov, I., Hairutdinova, A., Khafizova, F., Zakirova, E., Hafizov, R., and Kayumov, A. (2016). Soluble and immobilized papain and trypsin as destroyers of bacterial biofilms. Genes and Cells. 10, 106–112.

UniProt Consortium. (2014). UniProt: a hub for protein information. Nucleic acids Res. 43, D204–D212. doi: 10.1093/nar/gku989

Wang, S. L., Hsu, W. T., Liang, T. W., Yen, Y. H., and Wang, C. L. (2008). Purification and characterization of three novel keratinolytic metalloproteases produced by *Chryseobacterium indologenes* TKU014 in a shrimp shell powder medium. Bioresource Technol. 99, 5679–5686. doi: 10.1016/j.biortech.2007.10.024

Watnick, P., and Kolter, R. (2000). Biofilm, city of microbes. J. Bacteriol. 182, 2675–2679.

Weatherly, L. M., and Gosse, J.A. (2017). Triclosan exposure, transformation, and human health effects. J. Toxicol. Environ. Health. Part B. 20, 447–469. doi: 10.1080/10937404.2017.1399306

