## Supplementary file for "Enzymatic degradation of biofilm by metalloprotease from *Microbacterium* sp. SKS10"

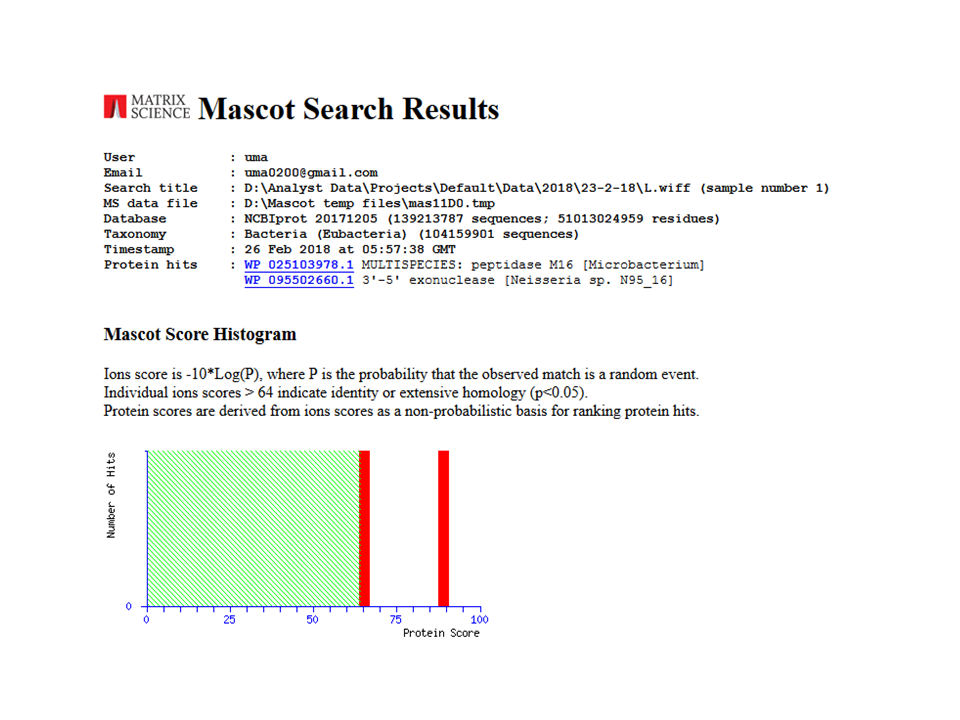

**Figure S1** Mascot search results

Table S1 Carbohydrate utilization tests of *Microbacterium* sp. SKS10

| Biochemical tests | |
| --- | --- |
| Positive | **Negative** |
| Glycerol utilization | Lactose utilization |
| Salicin utilization | Xylose utilization |
| Esculin hydrolysis | Maltose utilization |
| Citrate utilization | Fructose utilization |
| Malonate utilization | Dextrose utilization |
| Skim milk agar test | Galactose utilization |
|  | Raffinose utilization |
|  | Trehalose utilization |
|  | Dulcitol utilization |
|  | Sorbitol utilization |
|  | Mannitol utilization |
|  | Cellobiose utilization |
|  | Melizitose utilization |
|  | ɑ-methyl-D-mannoside utilization |
|  | Xylitol utilization |
|  | Melibiose utilization |
|  | Sucrose utilization |
|  | L-Arabinose utilization |
|  | Mannose utilization |
|  | Inulin utilization |
|  | Sodium gluconate utilization |
|  | Inositol utilization |
|  | Arabitol utilization |
|  | Erythritol utilization |
|  | ɑ-methyl-D-glucoside utilization |
|  | Rhamnose utilization |
|  | ortho-Nitrophenyl-β-galactoside utilization |
|  | D-Arabinose utilization |
|  | Sorbose utilization |

A single isolated bacterial colony of *Microbacterium* sp. SKS10 was inoculated in brain heart infusion broth and incubated at 35–37 °C till the OD reached 0.5 at 620 nm. 50 μl of this culture was inoculated in each well of HiCarbo kit followed by incubation at 35 ± 2 °C for 18–24 hours.

Table S2 Biochemical characterization of *Microbacterium* sp. SKS10

| Biochemical test | Interpretation | Result |
| --- | --- | --- |
| Methyl red test | Stable acids production | Negative |
| Vogues Proskauer test | Acetylmethylcarbinol production | Negative |
| Tryptone water test | Indole production | Negative |
| Starch agar test | Amylase production | Negative |
| Spirit blue agar test | Lipase production | Negative |
| Lysine decarboxylase test | Lysine decarboxylase production | Negative |
| Skim milk agar test | Protease production | Positive |

The ability of *Microbacterium* sp. SKS10 to produce stable acids, acetylmethylcarbinol, indole, amylase, lipase, lysine decarboxylase and protease was detected by inoculating single bacterial colony in respective media as mentioned in the text.
